# Hyperbolic geometry-based deep learning methods to produce population trees from genotype data

**DOI:** 10.1101/2022.03.28.484797

**Authors:** Aman Patel, Daniel Mas Montserrat, Carlos Bustamante, Alexander Ioannidis

## Abstract

The production of population-level trees using the genomic data of individuals is a fundamental task in the field of population genetics. Typically, these trees are produced using methods like hierarchical clustering, neighbor joining, or maximum likelihood. However, such methods are non-parametric: they require all data to be present at the time of tree formation, and the addition of new data points necessitates the regeneration of the entire tree, a potentially expensive process. They also do not easily integrate with larger workflows. In this study, we aim to address these problems by introducing parametric deep learning methods for tree formation from genotype data. Our models specifically create continuous representations of population trees in hyperbolic space, which has previously proven highly effective in embedding hierarchically structured data. We present two different architectures - a multi-layer perceptron (MLP) and a variational autoencoder (VAE) - and we analyze their performance using a variety of metrics along with comparisons to established tree-building methods. Both models tested produce embedding spaces that reflect human evolutionary history. In addition, we demonstrate the generalizability of these models by verifying that addition of new samples to an existing tree occurs in a semantically meaningful manner. Finally, we use Dasgupta’s Cost to compare the quality of trees generated by our models to those produced by established methods. Despite the fact that the benchmark methods are directly fit on the evaluation data, our models are able to outperform some of these and achieve highly comparable performance overall.

**Author summary:** Tree production is a vital task in population genetics, but current approaches fall prey to several common shortfalls. Most notably, they lack the ability to add new data points after tree generation, and they are often difficult to use in larger pipelines. By leveraging cutting-edge advances pairing deep learning with hyperbolic geometry, we develop multiple models designed to rectify these issues. Through experiments on a dataset of humans from globally widespread ancestries, we demonstrate the generalizability of our models to new data, and we also show strong empirical performance with respect to currently used methods. In addition, we show that the data representations produced by our models are semantically meaningful and reflect known facts about human evolutionary history. Finally, we discuss the additional benefits our models could provide, including improved visualization, greater privacy preservation, and improved integration with downstream machine learning tasks. In conclusion, we present models that are accurate, flexible, and generalizable, with the potential to facilitate a variety of further applications.

## Introduction

Recent advances in genetic sequencing technology, along with the wide availability of genetic data, have resulted in the increased interest in predicting disease risk and other traits directly from an individual’s genomic data. These predictions are made possible through studies like Genome-Wide Asssociation Studies (GWAS), which determine correlations between genomic variants and such traits, and the calculation of quantities like Polygenic Risk Scores (PRS), which use an individual’s genotype to quantify disease risk. However, these models do not always generalize across ancestries, often requiring different models to be developed for different populations [1]. Accurate knowledge of an individual’s genetic ancestry is therefore frequently important in making the most accurate and effective healthcare decisions possible.

Ancestry categories are hierarchical, with each individual having a label at each level in the hierarchy. For example, if we were to separate humans by continental population, a particular individual could be described as European. If we then proceeded to cluster the European population, the same individual could then be described as Italian. If we then identified sub-clusters within the Italian population, this person could now be identified as Sicilian, and so on. Trees are a natural choice to represent such hierarchically structured data, and they have been widely used in the past in several related paradigms. For example, Li et al. utilized trees from genotype data to elucidate migration patterns and study relationships between human populations from across the world [2]. Duda and Zrzavý combined genetic data with linguistic data to further probe these relationships [3]. Trees are widely applicable to non-human species as well, as evidenced by Parker et al.’s study utilizing trees to elucidate the population histories of canids [4]. On a broader scale, of course, trees have been utilized to catalog and categorize the entirety of life on Earth into an all-encompassing “tree of life” [5]. Finally, there has been much theoretical, mathematical, and computational work done to devise and improve state-of-the-art tree production methods [6–8].

Indeed, utilizing trees can provide several benefits in the study of ancestry. Importantly, a tree can help assess how related two population groups or two individuals are to each other. This fact can allow us to, for example, use population trees to determine if GWAS or PRS results from one group are likely to generalize to another. Similarly, we can use trees to determine which groups can be added to a particular study and therefore increase its power while retaining accuracy. At present, the majority of genome-trait association studies have been conducted in individuals of European descent, largely neglecting individuals of other ancestries [1]. Population trees can allow us to optimize study design for these groups and can be used for pharmacogenomics, thus allowing for the delivery of vital healthcare information as accurately and efficiently as possible [9].

Genetic ancestry-based trees also have a variety of applications outside healthcare. Notably, their structure can provide vital information regarding the course of human evolution, and they can also potentially explain various migration events that have occurred through history [2, 6]. Such information may prove valuable for historical or demographic studies. Furthermore, trees can be produced for crops, livestock, and other species, potentially aiding in decisions regarding agriculture and ecological health [4, 10].

To create a tree from genomic data, it is necessary to leverage certain genetic signatures to calculate measures of similarity between the entities that will comprise the tree. A common choice is to use Single Nucleotide Polymorphisms (SNPs), or single genomic positions that are known to vary across human populations due to the evolutionary history of the human species [2, 11]. In general, it is expected that a pair of closely related individuals will share more SNPs than a pair of more distantly related ones. Typically, a tree-building algorithm like hierarchical clustering is employed together with a similarity metric related to the number of SNPs shared in common between each pair of individuals [12]. However, such trees cannot be easily included as intermediate steps in larger workflows, and they are rigid and non-parametric - that is, it is not possible to introduce new samples into the tree without recreating the entire tree. This has two main disadvantages: first, it requires one to keep track of all the samples to recreate the tree. Second, it is highly computationally inefficient, and it is not a feasible solution for large modern biobanks and databases. Thanks to the decreasing cost of sequencing, more datasets and biobanks that include a large number of high-resolution SNP sequences are becoming available, generating a need for efficient and scalable alternatives to tree-generation techniques. Thus, a parametric method to embed tree nodes in a continuous space would, as illustrated in Fig 1, prove highly useful in the study of population genetics.

**Fig 1.**
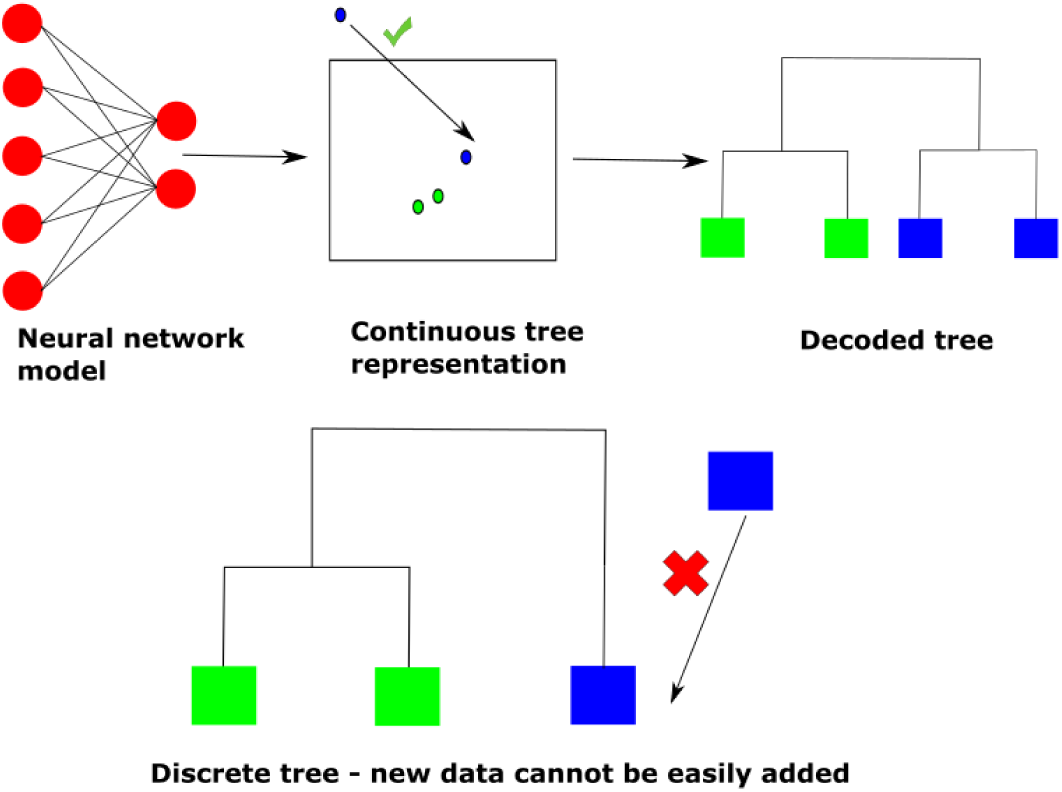
An advantage of using continuous tree representations rather than discrete trees is that new samples can be easily added to the current tree.

It has long been established that hyperbolic spaces are ideal for these embeddings, and a method called HypHC has recently been developed to embed data such that the embeddings reflect the structure of a hierarchical clustering tree [13]. Here, we present two parametric adaptations of HypHC: a multi-layer perceptron (MLP) model, and a Variational Autoencoder (VAE) model. We demonstrate the utility of these models to embed genotype data in an informative manner, and we compare the quality of the trees learnt to those produced by HypHC and hierarchical clustering. Overall, we demonstrate the utility of hyperbolic geometry in creating flexible parametric methods for tree creation for population genetics analysis.

## Methods

### Hyperbolic Space and Poincare Embeddings

Previous studies have demonstrated the great efficacy of hyperbolic spaces in learning embeddings for hierarchically structured data, including text, graphs, and in particular, trees [14–16]. We will therefore describe the properties of hyperbolic spaces - in particular, those following the Poincaré disc model - which allow for this hierarchical structure to be captured, along with those that are relevant for the understanding of our models.

In general, hyperbolic geometry is defined as a non-Euclidean geometry with a constant negative curvature [13, 17]. Various types of hyperbolic spaces exist, and one commonly used space is the Poincaré ball model, or its two-dimensional equivalent, the Poincaré disk. In a Poincaré disk (as with other hyperbolic spaces), the distance between two points *z*_*i*_ and *z*_*j*_ is calculated differently than in the standard Euclidean formulation [13]. Specifically, the distance formula is given by:

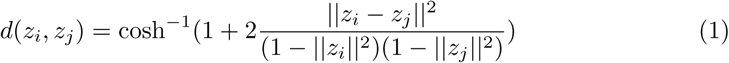

A byproduct of this distance metric is that surface area grows in an exponential manner as we reach the edge of the disk. Since the number of leaves in a tree grows exponentially with respect to the tree’s height, a Poincaré disk (or other hyperbolic space) is therefore ideal for learning accurate tree representations without the need for an excessively large embedding space. In fact, one can think of hyperbolic spaces as continuous analogues of trees [14]. Previous studies have demonstrated methods to embed trees into hyperbolic spaces with minimal error [16]. Euclidean spaces do not possess this property of exponential growth, meaning modeling larger trees requires a higher dimensionality, which can cause a variety of problems with training, including overfitting and memory concerns [14].

It is also important to consider the nature of the curves, or geodesics, which represent the shortest paths between two points in the Poincaré disk. In this formulation, geodesics are either lines through the origin or circular arcs that intersect the two points and are orthogonal to the edges of the disk [18].

Given our knowledge of geodesics, we must introduce one final important property: the *exponential map*. Assume that we start with a point *z*, which is on a unit geodesic *γ* such that *γ*(0) = *z*. There will exist only one possible *γ* such that the tangent vector at *z* (i.e. *γ′* (0)) is a certain vector *v*. We then define the exponential map of *v* as *exp*_*z*_(*v*) = *γ*(1). A closed-form solution exists for this transformation, and it is used frequently in the various architectures and probability distributions utilized in this study [17].

### The HypHC Model

Central to the models employed in this study is Hyperbolic Hierarchical Clustering, or HypHC, which was introduced by Chami et al. in 2020 [13]. This method utilizes a differentiable objective function to embed data points in hyperbolic space such that distances between embeddings reflect closeness in a hierarchical clustering tree. Hyperbolic geometry was chosen for this task because it allows for more natural and effective representations of trees than is possible in the standard Euclidean space [13]. Specifically, the objective function of HypHC is a continuous relaxation in hyperbolic space of Dasgupta’s Cost, a common metric used to evaluate the quality of hierarchical cluster assignments.

Given a similarity matrix *w* representing the data, and a tree *T* generated with hierarchical clustering, Dasgupta’s cost can be written as follows:

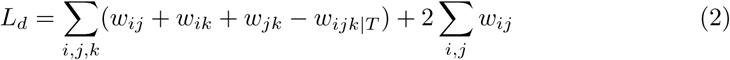

where *w*_*ij*_ is the similarity between sample *i* and *j, w*_*ijk*|*T*_ = *w*_*ij*_ if the tree first splits *k* from *i* and *j, w*_*ik*_ if the tree first splits *j* from *i* and *k*, and *w*_*jk*_ if the tree first splits *i* from *j* and *k* [13].

Dasgupta’s cost therefore calculates whether more similar nodes are located closer together in the tree than less related ones. Alternatively, it gauges if the Lowest Common Ancestors (LCA) - the points at which two nodes split from each other - of more similar nodes are lower in the tree, while the LCAs of less similar nodes are higher in the tree. The term *w*_*ijk*|*T*_ can therefore be interpreted as the similarity of the two nodes in {*i, j, k*} with the LCA that is farthest from the root of the tree [13].

Although this objective function is discrete, a continuous analog can be developed in hyperbolic space. To do so, we must define the continuous equivalent of the LCA. Let *z*_*i*_ and *z*_*j*_ be the hyperbolic embeddings of two data points *i* and *j*, and let *q*_*ij*_ be the curve representing the geodesic (shortest path distance) between *z*_*i*_ and *z*_*j*_. We first consider that in a conventional tree, the LCA between two nodes will be the point on the shortest path between the nodes that is closest to the root. Similarly, if we assume our tree is rooted at the origin in hyperbolic space, the hyperbolic LCA between *z*_*i*_ and *z*_*j*_ will be the point on the geodesic that is closest to the origin [13].

If we define *d*_*LCA*_(*z*_*i*_, *z*_*j*_) to be the distance from the origin to the hyperbolic LCA of *z*_*i*_ and *z*_*j*_, we can define the continuous analog to Dasgupta’s cost as follows:

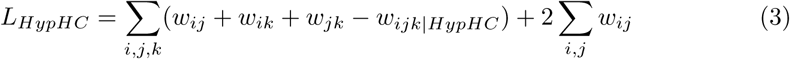

The term *w*_*ijk*|*HypHC*_ is then defined as follows:

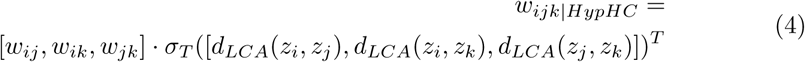

where *σ*_*T*_ is the softmax function with a learned temperature scaling parameter (specifically, the logits are scaled by 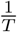 before the softmax is applied) [13]. Therefore, the softmax output acts as a smooth relaxation of an indicator function.

This objective can then be optimized by gradient descent in a similar manner to a conventional neural network. Even though all data points are considered at once, we can divide the triplets *i, j, k* into batches for the sake of computational efficiency and memory usage.

Finally, if desired, the HypHC paper provides algorithms to convert the continuous space into a discrete tree. The most theoretically rigorous of these can be computationally expensive under certain situations, but a greedy algorithm is provided which runs far faster and has been validated to achieve comparable performance. In this algorithm, points first are normalized to have the same distance from the origin in the Poincaré Disk. Next, the points are sorted according to their angles in the disk, which are analogous to their positions when normalized. We then calculate the two largest gaps in this sorted list (ie. the two largest differences between consecutive angles), and we split the dataset into points that have angles between these two gaps and points that do not. Now that the first division has been determined, we recursively divide each split at the largest gap until the full tree structure is reached [13]. In this manner, we have a simple, efficient algorithm to produce discrete trees from HypHC embedding spaces.

### Novel Parametric Models

We will now discuss the parametric models developed as part of this study.

#### Hyperbolic MLP Model

The HypHC framework can easily be extended to a parametric model, with the simplest example consisting of a fully connected multi-layer perceptron (MLP) with the HypHC loss as the loss function. Unlike the HypHC model, loss values are not calculated using all the training examples at once. Instead, we take the more conventional approach of dividing data into batches and applying gradient updates per batch. Specifically, for each individual in a batch, embeddings will be obtained using the fully connected layers. Next, the HypHC loss will be calculated using only these embeddings, and a gradient update will then be performed.

While the traditional HypHC model directly optimizes each embedding *z*_*i*_ given an input *x*_*i*_, our proposed Hyperbolic MLP model makes uses of a function *f* (represented by the neural network) to translate between input and output, specifically by calculating *f* (*x*_*i*_) = *z*_*i*_. Therefore, we learn a parametric model, *f*, that predicts *z*_*i*_ given an input *x*_*i*_.

When the model has been trained, embeddings for new data points can be obtained by applying the fully connected layers to the input genotype. In this manner, the embedding space is still optimized using a hierarchical clustering-based objective, but it also retains the ability for new data points to be added in a semantically meaningful manner.

#### Hyperbolic VAE Model

The model consists of MLP-based encoders and decoders, with transformations applied to convert data into coordinates in hyperbolic space, following the framework provided in [17]. Specifically, after applying linear layers, followed by ReLU non-linearities, to the input data to obtain predicted means and standard deviations for the embedding coordinates, the predicted means are applied along the exponential map (which does not occur in the MLP model). Latent coordinates are then sampled. The decoder consists of a logarithmic map - to undo the exponential map - followed by a series of linear layers to reproduce the input. In all cases, the embedding space used was two-dimensional.

The optimization objective of the model consists of three weighted components. The first two components result from the need to maximize the evidence lower bound (ELBO), as in standard VAEs. When extended to Riemannian spaces, this is equivalent to minimizing the expression [17]:

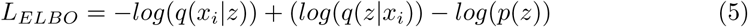

The term −*log*(*q*(*x*_*i*_ |*z*)) represents the likelihood of the reconstructed data, and it is calculated in practice using the binary cross-entropy between the reconstructions and the original data. The term (*log*(*q*(*z* |*x*_*i*_)) −*log*(*p*(*z*))) represents the similarity between the predicted and prior distributions for latent space coordinates, and it is analogous to the KL divergence term in the standard VAE objective. For the latent prior, in lieu of using a standard normal distribution, we use a Wrapped Normal distribution on a Poincare Ball manifold. The Wrapped Normal consists of a Normal Distribution mapped along the exponential map [17]. This term therefore consists of the difference in log probabilities between a Wrapped Normal distribution with each predicted mean and standard deviation, and a Wrapped Normal distribution with mean 0 and standard deviation 1.

The HypHC loss is the third and final component. This term is calculated batch-wise; the triplets used in the HypHC objective consist of all possible combinations of three data points from the current batch.

The final per-batch objective for the model can therefore be written as:

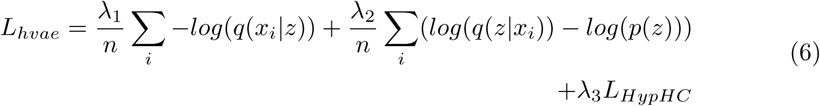

where *λ*_1_, *λ*_2_, and *λ*_3_ are manually specified weights for each of the loss terms and *n* is the batch size. In this architecture, the reconstruction loss is intended as an additional objective to ensure the latent embedding space most accurately reflects the relationships present in the raw data. We surmised that the combination of HypHC loss and reconstruction loss could possibly prove more effective than simply using the HypHC loss alone.

Furthermore, the model’s decoder could allow for several other promising applications, including the reconstruction of genotype data from the embedding space and the production of synthetic data with specified properties. However, as this study focuses on tree creation, these properties are not extensively tested and benchmarked and are left as future work.

### Dataset and Training

For all experiments, data was derived from the 1000 genomes project, the Human Genome Diversity Project (HGDP), and the Simons Genome Diversity Project (SGDP) [19–21]. The combined dataset consists of genotype data at 23,155,158 SNPs for a total of 2,965 individuals, or 5,930 haploid genotypes. Individuals are derived from a total of 136 different ethnic groups, which in turn are grouped into 7 continental regions: Africa, Europe, the Americas, Oceania, West Asia, East Asia, and South Asia. For the purposes of this study, each haploid genotype was considered a separate data point, and these genotypes were randomly divided into training, validation, and test data at a ratio of 80% to 10% to 10%. The input feature set was then filtered to only keep the 500,000 SNPs with the highest variance on a random sample of 2,000 genotypes from the training data. We then trained the MLP and the VAE models on this data, using a manual search to find effective hyperparameters. For the hyperbolic aspects of the parametric models (eg. exponential map, Wrapped Normal distribution, Poincaré Disk), we utilized implementations found in the Poincare VAE repo by Mathieu et al., and our implementation of the HypHC loss was adapted from the HypHC repo [13, 17]. Notably, to ensure effective visualization, only two-dimensional embedding spaces were used.

We conducted a non-exhaustive hyperparameter and architecture search for each model, which allowed us to arrive at the following values. The MLP model consisted of three hidden layers, with sizes of 300, 200, and 100, and ReLU activation and dropout after each layer. The encoder of the VAE model consisted of the same layer sizes, with batch normalization in addition to ReLU and dropout with probability 0.2. The decoder of the VAE model consisted of hidden layers of size 100, 200, and 300. All embeddings were two-dimensional. Schematics of the models are illustrated in Fig 2

**Fig 2.**
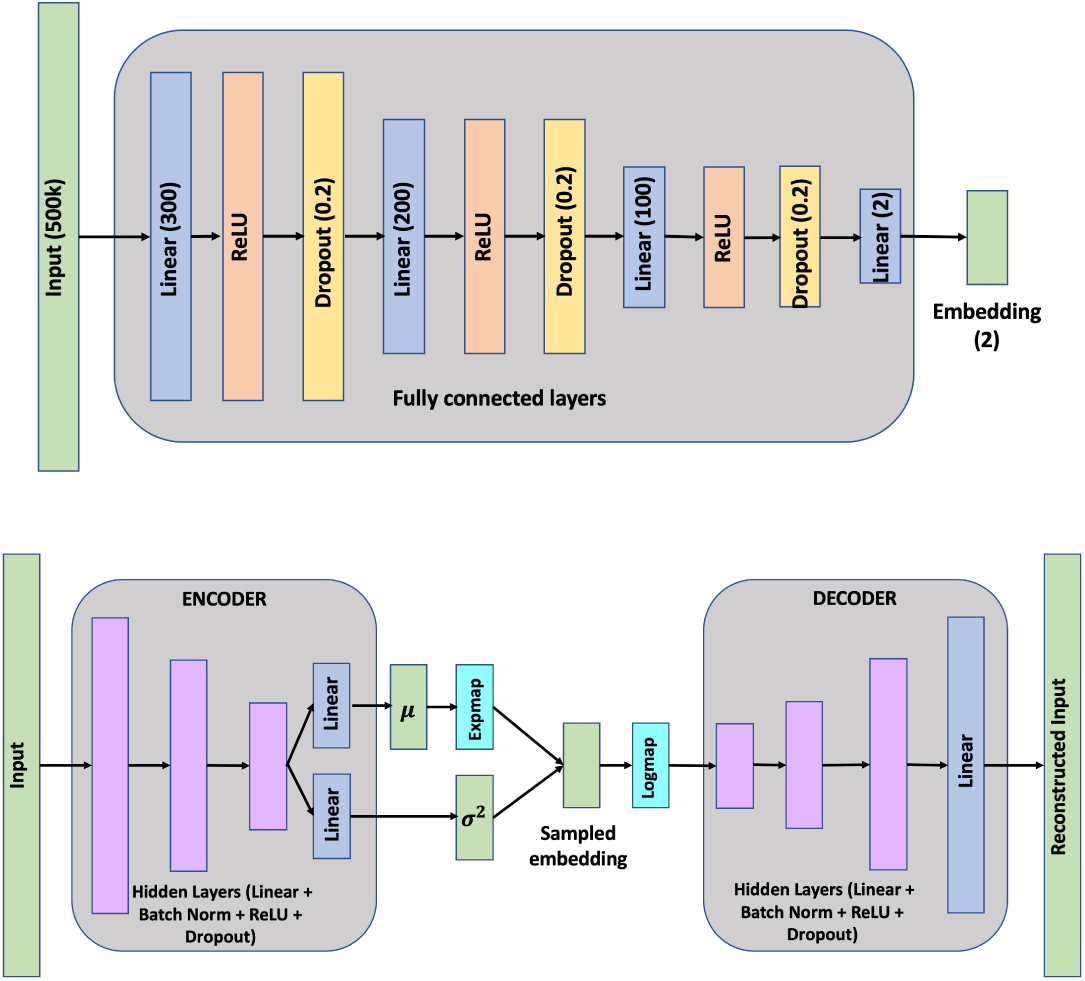
Schematics of the MLP (top) and VAE (bottom) models utilized in training. All layer sizes in the VAE match those in the MLP. Green boxes represent data or values, and all other colors represent operations.

The MLP model was trained with a learning rate of 10^−3^ and a temperature of 10^−4^, and the VAE model was trained with a learning rate of 10^−2^ and a temperature of 10^−6^. Due to the nature of the HypHC loss calculation, the batch size is more consequential than in most machine learning models. In both cases, a batch size of 64 was used to balance runtime with a sufficient sample size for HypHC loss. In all cases, the Riemannian Adam optimizer was used [13, 22]. Finally, all models were trained using early stopping with a patience of 5.

After inspecting the embedding spaces that were learnt, we compared the results of these models to those from non-parametric models applied to the test dataset – namely, HypHC and various forms of hierarchical clustering. In all cases, the similarity between two data points was defined as the fraction of SNPs for which the same nucleotide was present.

## Results

### Structure of Embedding Space

It is first instructive to determine whether the tree embedding space contains informative representations which reflect relationships between ancestries. Fig 3 displays the continuous tree representations produced by selected VAE (top) and MLP (bottom) models when evaluated on the test dataset. In both cases, most continental populations separate into clear clusters, thus accurately capturing the genetic differences between these groups. Furthermore, the ordering and spacing of these clusters largely reflects the predominant hypotheses of human migration and peopling of the planet. According to these theories, the earliest *Homo sapiens* populations were found in Africa, and over thousands of years, certain groups crossed into Eurasia and subsequently reached all corners of the Earth. Thus, all non-African humans arose from a limited number of founders who left Africa, meaning they share greater genetic similarity with each other than they do with the more diverse populations of Africa. The spacing of clusters in both plots - namely, the fact that the African cluster is the most separated from the others - therefore perfectly represents established scholarship [23, 24].

**Fig 3.**
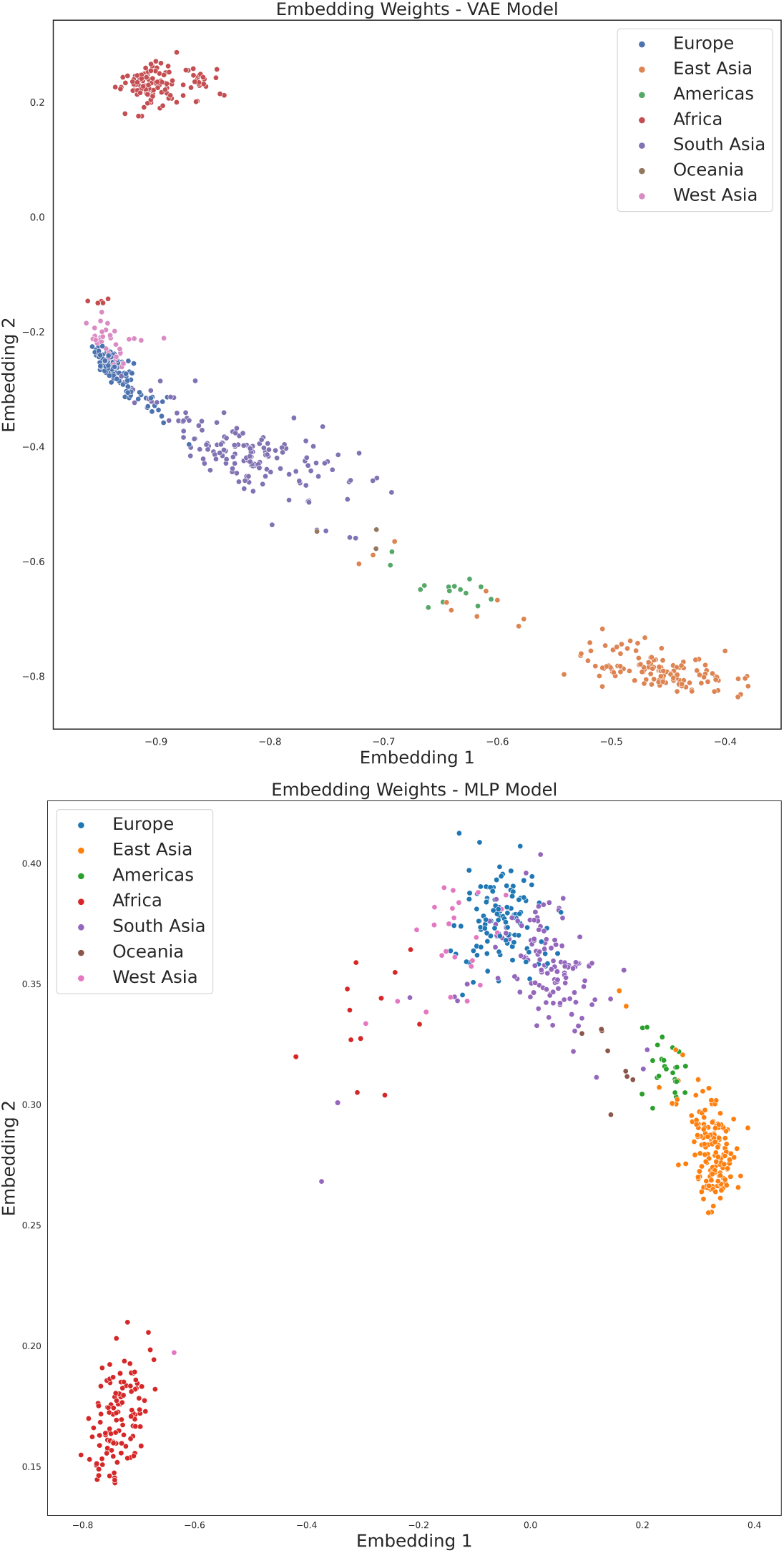
Structure of embedding spaces produced on the test set by the VAE (top) and MLP (bottom) models.

Similarly, the order of continental populations in the plot reflects the history of human migration. When early human populations left Africa, they likely did so via the modern near East, which falls under the “West Asia” category in our dataset. From there, migrations are believed to have occurred to the east into Southern Asia, and eventually to the west into Europe. Within Asia, migrations occurred to populate the entire continent, and along the Indian Ocean coast into Oceania [25, 26]. Finally, humans crossed the Bering Strait land bridge from East Asia into the Americas. Due to West Asia being the immediate destination from Africa, and a region that has remained in contact with it, it is reasonable that West Asians are the closest groups to Africans in the plot. It is also fitting that Europeans and South Asians are closest to West Asians, due again both to geography and the migration patterns mentioned above. Finally, due to the patterns of migration within and out of Asia, we would expect South Asians, East Asians, Oceanians, and Americans to be present in close proximity to each other, which is indeed the case.

The model can also deliver insights regarding the unique genetic ancestries of specific ethnic groups. For example, consider the African (red) points that are located nearby to the West Asian (pink) population. These individuals belong to either the Mozabite or Saharawi ethnic groups. The Mozabites are a small Berber ethnic group in Algeria, while the Saharawi occupy the disputed territory of Western Sahara. Due to their locations in North Africa, both groups have strong cultural, linguistic - and genetic - influences from the Arabian Peninsula. Thus, their placement between the West Asians and the rest of the African populations is an accurate reflection of their history [27, 28]. In conclusion, both the MLP and VAE models produce embedding spaces that largely reflect established knowledge regarding the genetic relationships between human ancestries.

### Insertion of New Genotypes

As stated, the chief advantage of parametric tree production methods is that new samples can be added to an existing tree in a semantically meaningful manner. We therefore performed a simple experiment to verify this functionality holds for our models. First, we produced embeddings for all training examples, thus generating the tree represenation for the entire training dataset. Next, we randomly selected 20 examples from the test dataset and added those to the embedding space. As evidenced in Fig 4, the test examples fall within their respective continental populations. Even small populations like Oceanians, who occupy a limited area in the training embedding space and therefore may not generalize well to unseen samples, display effective inference.

**Fig 4.**
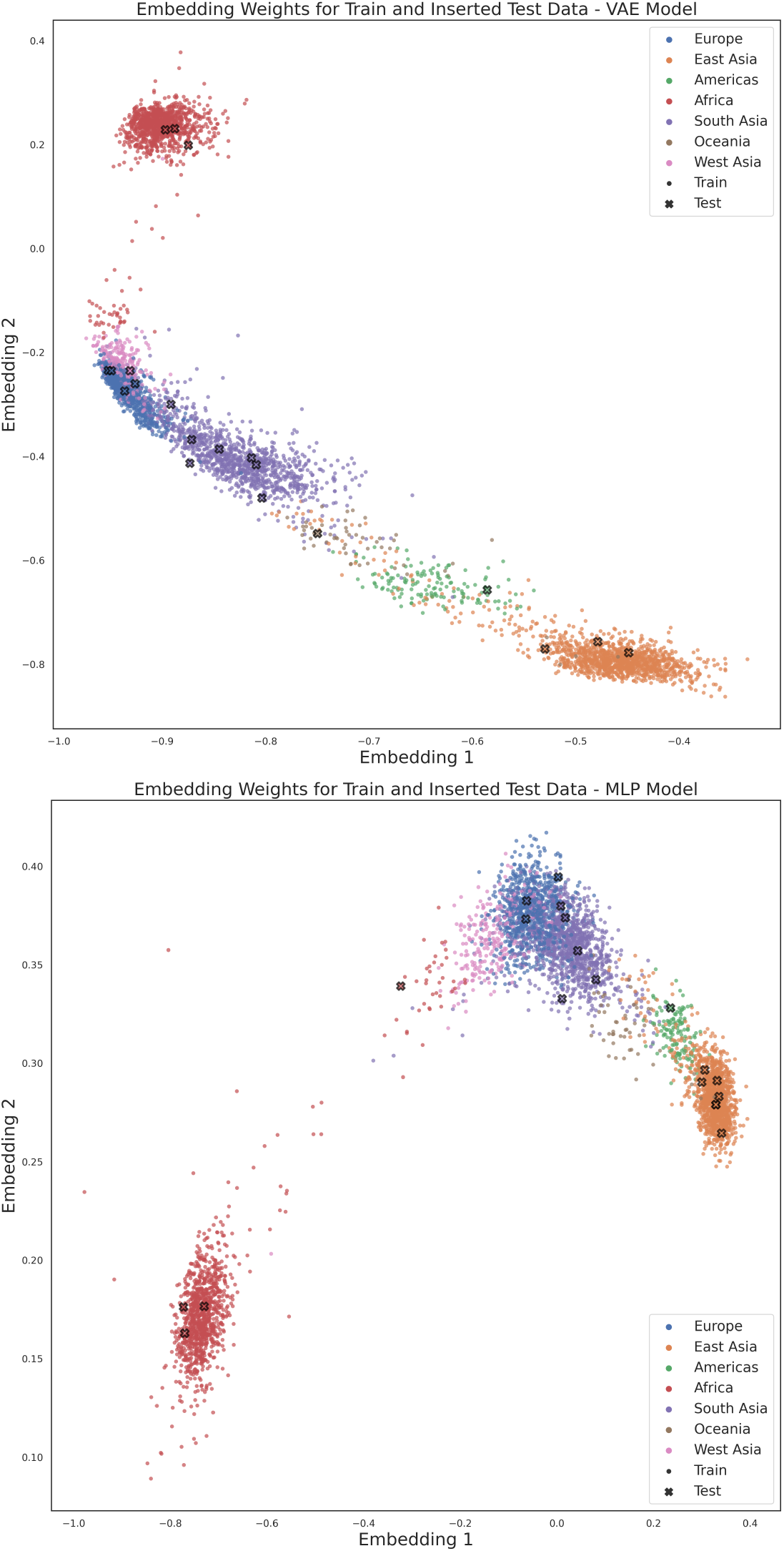
Outcome of adding randomly selected test samples to a tree formed from the training set by the VAE (top) and MLP (bottom) models. Training samples are represented by dots, and test samples are represented by crosses with black edges.

These results therefore illustrate that the models presented in this study are able to effectively generalize beyond the data they were trained on. Specifically, one can easily incorporate new samples into existing tree representations, a useful task for a variety of experimental settings.

### Performance Comparison

To explore comparisons with non-parametric methods, we performed the following procedure for the MLP and VAE models. First, we used a trained model to predict on all samples from the test dataset. These embeddings represent a continuous tree consisting of all test datapoints, and we then converted them into a discrete tree using the decoding method suggested in the HypHC paper. We then compared this tree to a variety of non-parametric methods, with Dasgupta’s cost as the metric of choice. As stated, these non-parametric methods include HypHC and various forms of hierarchical clustering. These variations of hierarchical clustering differ in the manner in which distances between clusters are calculated. If we assume we are calculating the distances between clusters *u* and *v*, then these methods can be described as follows:

- **Nearest Point Algorithm:** *d*(*u, v*) = *min*(*d*(*u*[*i*], *v*[*j*])
- **Farthest Point Algorithm:** *d*(*u, v*) = *max*(*d*(*u*[*i*], *v*[*j*]))
- **UPGMA Algorithm:** 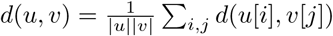
- **WPGMA Algorithm:** 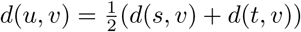 where cluster *u* was created from clusters *s* and *t*.

These methods represent all hierarchical clustering variations found in the Scipy Python package able to utilize a Hamming Distance metric [29]. HypHC was implemented by adapting code found in the HypHC package to fit the specific use case [13]. For the HypHC model, we used a learning rate of 10^−2^, a temperature of 10^−3^, and early stopping with a patience of 5.

Note that since the output of the VAE model is non-deterministic, the value presented is the result of aggregating the Dasgupta’s Cost values produced by predicting on the test set ten different times. Table 1 shows the Dasgupta’s Cost values for all methods on the test set.

**Table 1.**
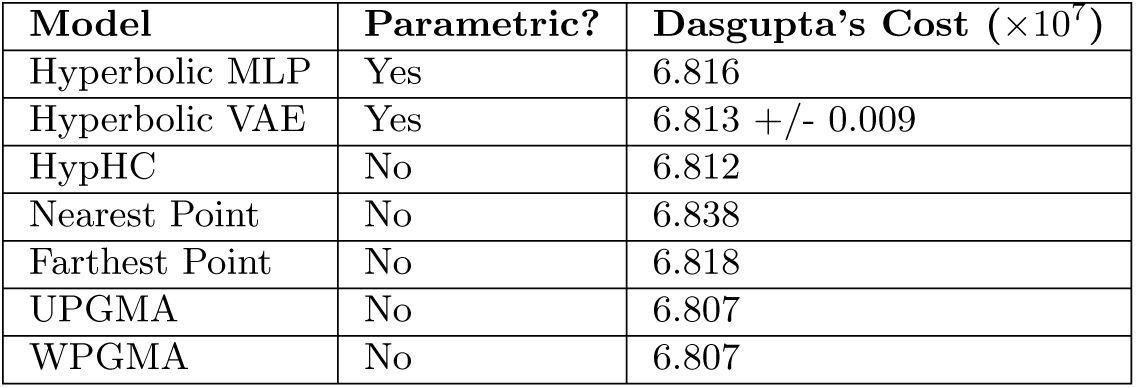
Comparison of Dasgupta’s Cost values on the test dataset for all parametric and non-parametric methods tested. Since the output of the VAE model is non-deterministic, the value presented is the result of predicting on the test set ten different times.

| Model | Parametric? | Dasgupta’s Cost ($\times 10^7$) |
| --- | --- | --- |
| Hyperbolic MLP | Yes | 6.816 |
| Hyperbolic VAE | Yes | 6.813 +/- 0.009 |
| HypHC | No | 6.812 |
| Nearest Point | No | 6.838 |
| Farthest Point | No | 6.818 |
| UPGMA | No | 6.807 |
| WPGMA | No | 6.807 |

As shown in Table 1, both parametric methods achieved highly competitive performance as compared to the non-parametric methods, even outperforming multiple of these benchmarks. It is instructive to once again note the key difference between the parametric and non-parametric methods with respect to this experiment - while the test set was directly used to train or fit all non-parametric methods used in this experiment, it represents completely unseen data for the parametric methods. The fact that the VAE and MLP models performed similarly to the state-of-the-art methods despite this disadvantage reflects their generalizability and ability to create effective tree representations from previously unseen sequences.

Notably, the performances of the VAE and MLP models were within the margin of error from each other, indicating the efficacy of both training objectives in the creation of continuous tree representations. Further experimentation is possibly needed to determine the optimal strategy.

## Discussion

Currently, if one desires to create a population tree from individual genotype data, standard tree building algorithms require all data to be present at the time of tree creation. If new data is obtained subsequently, it cannot easily be added to the resultant tree; instead, the tree must be remade. In addition, as the method is discrete, it cannot easily be incorporated into larger workflows. An effective parametric analog for hierarchical clustering that can successfully represent genotype data is therefore desirable. Hyperbolic geometry is generally seen as a natural choice for embedding hierarchically structured data, so we focused in this area - and in particular, on variations of the HypHC method - when designing methodologies for this study.

Overall, the performance of our two parametric methods - VAE and MLP-based models - proved very promising when evaluated from a variety of different perspectives. We first verified that our models’ embedding spaces grouped individuals by their continental population, and we observed that the location and orientation of these clusters reflected human evolutionary history. We also found cases in which subpopulations with diverse influences were represented accordingly. We next tested whether, given a tree already embedded into the model’s latent space, new samples were projected in a semantically meaningful manner. Finally, when comparing Dasgupta’s cost values on the test dataset, both the VAE model and MLP model fell solidly in the midst of the various hierarchical clustering methods tested. Taken together, these results allow us to conclude that both models tested produce effective tree representations and generalize well to unseen samples.

The properties of these models can allow for a variety of useful and interesting applications. Of course, one of the most obvious pertains to adding a new sample to a tree with a large number of nodes. Instead of performing the computationally expensive process of remaking the tree each time a new node is added, we can simply add points to the embedding space as we desire. A discrete tree can then be decoded from the embedding space. In this manner, significant time is saved when producing updated trees. Additionally, the frameworks presented makes it far easier to include hierarchical clustering as part of larger machine learning workflows. Embeddings can be calculated at some point in a pipeline, fed into the next step, and the entire model can eventually be trained in an end-to-end fashion. One can imagine, for example, using hyperbolic embeddings as inputs to a classifier for whether individuals are at risk for a particular disease. Ultimately, a continuous and parametric model can greatly expand the realm of tasks over which hierarchical clustering can contribute to a solution. Our models can also potentially aid in preserving privacy of genetic data. If researchers desire to produce a tree from certain individuals, they would normally need to access the raw genotype data. Now, another option could be to receive the model embeddings instead, after which the tree can be easily decoded. In this manner, the raw data will remain hidden, thus protecting the privacy of the subjects. Furthermore, researchers without permission to view the data could still perform analyses. Of course, such models would likely require modifications, along with vigorous testing, to ensure they are truly privacy-preserving. However, the models presented in this study represent a promising approach. Finally, the dual methods of representation possible - a continuous embedding space or a discrete tree - can allow for more comprehensive visualization of a population. A tree is clearly effective at denoting the position of each node in the overall hierarchy. However, relative distances between large groups or overall population trends may be more intuitively depicted in a continuous plot.

It is also informative to compare the two parametric models. The MLP and VAE achieved similar performance to each other, indicating that neither objective function is intrinsically superior to the other. This is an interesting finding, as the MLP objective is explicitly optimized for hierarchical clustering, while the VAE objective is not. It is possible that for this data type simply embedding the data in hyperbolic space - which, as stated previously, is highly effective for representing hierarchically structured data - is enough to achieve strong hierarchical clustering performance. Both objective functions can certainly produce semantically meaningful embeddings. Further elucidating the specific behavior imparted by each objective function is an interesting future direction, as is probing a larger number of architectures and objectives to determine the optimal design choices. Another area of exploration pertains to fully studying the capabilities of the VAE model. VAEs have a variety of functions that were not explicitly tested and benchmarked in this study, including reconstruction of input data and generation of new samples with desired properties [30]. Both properties could prove highly useful for genetic ancestry. For example, to observe the genetic characteristics of particular ethnicities, one could generate synthetic samples from desired locations in the embedding space and analyze the resulting genotypes. By observing which SNPs in the reconstructed genomes remain constant and which are allowed to vary, one can more deeply understand the characteristic genetic signatures of the ethnic group in question. Similar experiments can be performed for individuals of unknown origin. A VAE can be used to produce several reconstructions of the individual’s genotype, and by analyzing the variation present in these reconstructions, one can more fully discern the characteristics of the individual’s source population. A multifaceted model that is effective in each of these areas, while still producing an embedding space optimized for hierarchical clustering, would be a useful tool for the study of population genetics.

Several other areas of future work also exist. Most notably, while the models separated continental populations from each other in the embedding space, stratification of individual ancestries within continents was not nearly as effective. The most likely explanation for this issue is simply a lack of training data. To improve performance further, one would likely have to obtain more samples for each population or generate them synthetically. More data would also allow us to train targeted models on specific populations rather than individuals from around the world, thus allowing for greater regional precision and resolution. One could also experiment with the metrics used for hierarchical clustering. In all experiments performed in this study, the metric used was directly derived from the genotype data, which was also the input to the model. However, the similarity metric could also conceivably be used to add new information. For example, one could craft a similarity metric around whether two samples are of the same ethnicity, thus introducing supervised learning based on a sample’s population labels. One could then take this idea a step further and introduce arbitrary similarity metrics - for example, defining the metric based on the primary language spoken by each individual. If a model can cluster according to this metric, while still grouping by genotype within each cluster, it would prove an interesting tool.

In summary, this study introduces multiple parametric methods for tree formation from genotype data, and it also puts forward several future directions for portable private models, supervised approaches, and synthetic data generation based on these methods.

## Acknowledgments

The authors would like to thank Dr. Guhan Venkataraman for assistance with the preparation some of the datasets used in this study.

## Code Availability

Code for this study can be found at https://github.com/AI-sandbox/hyperLAI

## References

1. Martin AR, Kanai M, Kamatani Y, Okada Y, Neale BM, Daly MJ. Clinical use of current polygenic risk scores may exacerbate health disparities. Nature Genetics. 2019;51(4):584–591. doi:10.1038/s41588-019-0379-x.

2. Li JZ, Absher DM, Tang H, Southwick AM, Casto AM, Ramachandran S, et al. Worldwide human relationships inferred from genome-wide patterns of variation. Science. 2008;319(5866):1100–1104. doi:10.1126/science.1153717.

3. Duda P, Zrzavý J. Human population history revealed by a supertree approach. Scientific Reports. 2016;6(1). doi:10.1038/srep29890.

4. Parker HG, Dreger DL, Rimbault M, Davis BW, Mullen AB, Carpintero-Ramirez G, et al. Genomic analyses reveal the influence of geographic origin, migration, and hybridization on modern dog breed development. Cell Reports. 2017;19(4):697–708. doi:10.1016/j.celrep.2017.03.079.

5. Hug LA, Baker BJ, Anantharaman K, Brown CT, Probst AJ, Castelle CJ, et al. A new view of the Tree of Life. Nature Microbiology. 2016;1(5). doi:10.1038/nmicrobiol.2016.48.

6. Cavalli-Sforza LL, Edwards AW. Phylogenetic analysis: Models and estimation procedures. Evolution. 1967;21(3):550. doi:10.2307/2406616.

7. Felsenstein J. Evolutionary trees from gene frequencies and quantitative characters: Finding maximum likelihood estimates. Evolution. 1981;35(6):1229. doi:10.2307/2408134.

8. Pickrell JK, Pritchard JK. Inference of population splits and mixtures from genome-wide allele frequency data. PLoS Genetics. 2012;8(11). doi:10.1371/journal.pgen.1002967.

9. Huddart R, Fohner AE, Whirl-Carrillo M, Wojcik GL, Gignoux CR, Popejoy AB, et al. Standardized biogeographic grouping system for annotating populations in pharmacogenetic research. Clinical Pharmacology & Therapeutics. 2019;105(5):1256–1262.

10. Saslis-Lagoudakis CH, Savolainen V, Williamson EM, Forest F, Wagstaff SJ, Baral SR, et al. Phylogenies reveal predictive power of traditional medicine in bioprospecting. Proceedings of the National Academy of Sciences. 2012;109(39):15835–15840. doi:10.1073/pnas.1202242109.

11. DeGiorgio M, Jakobsson M, Rosenberg NA. Explaining worldwide patterns of human genetic variation using a coalescent-based serial founder model of migration outward from Africa. Proceedings of the National Academy of Sciences. 2009;106(38):16057–16062. doi:10.1073/pnas.0903341106.

12. Bouaziz M, Paccard C, Guedj M, Ambroise C. Ships: Spectral hierarchical clustering for the inference of population structure in genetic studies. PLoS ONE. 2012;7(10). doi:10.1371/journal.pone.0045685.

13. Chami I, Gu A, Chatziafratis V, Ré C. From Trees to Continuous Embeddings and Back: Hyperbolic Hierarchical Clustering; 2020.

14. Nickel M, Kiela D. Poincaré Embeddings for Learning Hierarchical Representations. In: Guyon I, Luxburg UV, Bengio S, Wallach H, Fergus R, Vishwanathan S, et al., editors. Advances in Neural Information Processing Systems. vol. 30. Curran Associates, Inc.; 2017.Available from: https://proceedings.neurips.cc/paper/2017/file/59dfa2df42d9e3d41f5b02bfc32229dd-Paper.pdf.

15. Tifrea A, Bécigneul G, Ganea OE. Poincaré GloVe: Hyperbolic Word Embeddings; 2018.

16. Sarkar R. Low distortion delaunay embedding of trees in hyperbolic plane. Graph Drawing. 2012; p. 355–366. doi:10.1007/978-3-642-25878-734.

17. Mathieu E, Lan CL, Maddison CJ, Tomioka R, Teh YW. Continuous Hierarchical Representations with Poincaré Variational Auto-Encoders; 2019.

18. Brannan DA, Esplen DA, Gray J. Geometry. Cambridge University Press; 2012.

19. Cavalli-Sforza LL. The Human Genome Diversity Project: Past, Present and Future. Nature Reviews Genetics. 2005;6(4):333–340. doi:10.1038/nrg1579.

20. Mallick S, Li H, Lipson M, Mathieson I, Gymrek M, Racimo F, et al. The Simons Genome Diversity Project: 300 genomes from 142 diverse populations. Nature. 2016;538(7624):201–206. doi:10.1038/nature18964.

21. Auton A, Abecasis GR, Altshuler DM, Durbin RM, Abecasis GR, Bentley DR, et al. A global reference for human genetic variation. Nature. 2015;526(7571):68–74. doi:10.1038/nature15393.

22. Bécigneul G, Ganea OE. Riemannian Adaptive Optimization Methods; 2019.

23. Tishkoff SA, Reed FA, Friedlaender FR, Ehret C, Ranciaro A, Froment A, et al. The genetic structure and history of Africans and African Americans. Science. 2009;324(5930):1035–1044. doi:10.1126/science.1172257.

24. Campbell MC, Tishkoff SA. African genetic diversity: Implications for human demographic history, modern human origins, and complex disease mapping. Annual Review of Genomics and Human Genetics. 2008;9(1):403–433. doi:10.1146/annurev.genom.9.081307.164258.

25. deMenocal PB, Stringer C. Climate and the peopling of the world. Nature. 2016;538(7623):49–50. doi:10.1038/nature19471.

26. Bae CJ, Douka K, Petraglia MD. On the origin of modern humans: Asian perspectives. Science. 2017;358(6368). doi:10.1126/science.aai9067.

27. Encyclopaedia Brittanica Eo. M’Zabite; 2015. Available from: https://www.britannica.com/topic/Mzabite.

28. Encyclopaedia Brittanica Eo. Western Sahara; 2021. Available from:https://www.britannica.com/place/Western-Sahara#ref14210.

29. Scipy.cluster.hierarchy.linkage, https://docs.scipy.org/doc/scipy/reference/generated/scipy.cluster.hierarchy.linkage. Available from: https://docs.scipy.org/doc/scipy/reference/generated/scipy.cluster.hierarchy.linkage.html.

30. Doersch C. Tutorial on Variational Autoencoders; 2021.

